# Turning green in microgravity: facial color changes as best physiological indicator of space sickness

**DOI:** 10.64898/2026.03.06.708504

**Authors:** Tess Bonnard, Emilie Doat, Jean-René Cazalets, Clément Morgat, Dominique Guehl, Etienne Guillaud

## Abstract

Motion sickness (MS) is commonly hypothesized to arise from sensory conflicts between incongruent sources of sensory information. Different types of sensory conflicts can induce MS, yet it remains unclear whether distinct contexts produce different physiological responses. Moreover, there is a lack of reliable objective predictors of MS, particularly for space motion sickness (SMS), which appears unrelated to motion sickness susceptibility on Earth. This study examined multiple physiological measures as potential objective markers of MS, including heart rate, blood pressure, salivary cortisol, skin conductance, skin surface temperature, and facial skin colorimetry. Subjective motion sickness severity and symptomatology were assessed using standardized questionnaires (SSQ, MSAQ, MSSQ). All measures were collected before and immediately after exposure to two sensory conflict paradigms: virtual reality (visuo-vestibular conflict) and parabolic flight (otolitho-canal conflict). Post-exposure, both paradigms were associated with increased cortisol, skin conductance, and skin greeness. Notably, increased skin greenness was associated with greater MS severity in parabolic flight and strongly correlated with subjective nausea ratings in both paradigms. Skin temperature and systolic blood were affected differently by VR and parabolic flight. No robust new physiological predictors of MS were identified. Overall, our findings suggest that facial skin color -particularly skin greenness- may serve as a simple, non-invasive, and reliable objective indicator of MS severity.

## INTRODUCTION

Motion sickness (MS) is a widespread condition that arises when individuals are exposed to unusual motion environments, leading to a range of unpleasant subjective sensations and physiological responses. Its origins span traditional forms—such as car, sea, and air sickness—to newer variants like cybersickness and space sickness, which occur in virtual environments and weightlessness settings. MS impair both motor and cognitive performance, leading to declines in short-term memory (Dahlman et al., 2009), decision- making, visual processing, concentration, reasoning, psychomotor and sensorimotor skills (Tal et al., 2012; Metzulat et al., 2025). This is particularly critical in operational contexts such as spacecraft landings or drone piloting, where high levels of cognitive and motor precision are required (Clément, 2020).

The most widely accepted explanation is the sensory conflict theory, which posits that motion sickness arises when sensory inputs from different modalities (e.g., visual and vestibular) are incongruent, generating a conflict in multi-sensory integration that triggers the associated symptoms (Reason & Brand, 1975). According to this theory, motion sickness can be categorized into two main types: inter-sensory conflict, as seen in classical motion sickness and cybersickness, and intra-sensory conflict, as intra vestibular (otolitho-canalar) conflict hypothesized in spaceflight (Gallagher et al., 2018). Differences in motion sickness susceptibility across contexts reflect the varying degrees and types of sensory conflict induced by different environments. The prevalence of motion sickness is approximately 50% among car passengers (Schmidt et al., 2020), 1% among airline passengers (Leung et al., 2019), and 60–80% among astronauts during the first few days of a mission (Heer et al., 2006). Motion sickness affects about 27% of astronauts upon return from short-duration flights (Ortega & Harm, 2008) and up to 100% following long-duration missions (Reschke et al., 2017c). Early experimental work highlighted difference in individual susceptibility among the context: Lentz (1984) reported very low or non- existent correlations between susceptibilities measured across laboratory paradigms such as Coriolis stimulation, visuo-vestibular conflict, and off-vertical rotation. Similarly, susceptibility measured on Earth failed to predict space motion sickness in Skylab crewmembers (Homick, 1979; Clément et al., 2023), and to predict vomiting incidence in space-like conditions such as parabolic flight (Golding et al., 2017). This emphasize that exposure to altered gravitational environments introduces novel sensory conditions not captured by terrestrial tests. Only weak to moderate correlations between earth MS susceptibility and cybersickness were observed (Kourtesis et al., 2023; Séba et al., 2023). Nevertheless there is a pressing need to predict, from objective measures, how an individual will respond to a novel inertial environment and whether they will develop motion sickness.

The Motion Sickness Susceptibility Questionnaire (MSSQ) provides a standardized and validated method for predicting individual vulnerability to motion sickness (Golding et al., 2006). As a self-report tool, the MSSQ relies on subjective recall, which can introduce reporting bias and limits. Objective monitoring of motion sickness focuses on symptoms such as sweating, fluctuations in body temperature, pallor, hypersalivation, nausea, and, in severe cases, vomiting (Zhang et al., 2016). These symptoms reflect activation of the autonomic nervous system, primarily mediated through specific brainstem regions such as the area postrema (chemoreceptor trigger zone), vagal pathways (Hasler, 2013; Oman & Cullen, 2014), and vestibular afferents (Yates et al., 2014; Cohen et al., 2019). The resulting physiological responses have been widely monitored in previous studies (Dobie, 2019 for review). For instance, both phasic and tonic components of skin conductance have been shown to correlate with motion sickness severity (Wan et al., 2003; Daviaux et al., 2019). Individuals suffering from motion sickness also exhibit change in blood pressure and heart rate (LaCount et al., 2009; Javaid et al., 2018), elevated cortisol levels (Otto et al., 2006; Choukèr et al., 2008), and reductions in core body temperature (Nobel et al., 2012; Nalivaiko, 2018).

In visually induced motion-sickness, facial skin temperature showed one of the strongest associations with severity (Keshavarz et al., 2022), and frequency-domain heart-rate variability measures were linked to levels of discomfort (Lobo et al., 2022). Interestingly, pupil dilation emerged as a predictor of cybersickness (Kourtesis et al., 2023). However, most of this research concerns visually induced sensory conflict; comparatively few studies have characterized physiological trajectories during parabolic flight or other space-analog environments and related them to individual susceptibility. Those investigations that do exist report limited or inconsistent predictive value for baseline physiological markers (Homick, 1978; Harm & Schlegel, 2002; Golding et al., 2017).

The aim of this study was to examine the evolution of physiological parameters during exposure to two distinct types of sensory conflict: a visuo-vestibular conflict induced by virtual reality (VR) and a vestibulo-vestibular conflict experienced during parabolic flight (PF). We sought to determine whether these two paradigms would elicit different physiological responses and whether the physiological changes observed in one paradigm would correlate with those observed in the other. Furthermore, by considering baseline pre-exposure values, we aimed to assess whether any baseline physiological parameter could serve as a predictor of individual susceptibility to motion sickness across both environments. To this end, we measured a broad set of physiological parameters, including blood pressure, heart rate, skin conductance, skin color, salivary cortisol, and skin surface temperature, at multiple time points before, during, and after exposure to virtual reality and parabolic flight.

### MATERIAL & METHODS

In this study, a single group of 29 healthy volunteers (mean age = 27 ± 7 years; 14 women, 15 men) was exposed to two distinct motion-sickness–inducing conditions: a visuo- vestibular conflict causing cybersickness, and gravitational variations during parabolic flight. 21 participants took part both to parabolic flight (PF) and virtual reality (VR) (27 ± 7 years; 9 women, 12 men). Additionally, 2 participants completed only the PF protocol (38 ± 11 years; 1 woman, 1 man) and 6 participants completed only the VR protocol (22 ± 2 years; 4 women, 2 men). For participants enrolled in both protocols, the order of the PF and VR sessions was counterbalanced, with a one-month interval between experiments. None of the participants reported any health conditions or prior medical procedures that could have affected their physiological responses. All individuals selected for the PF protocol completed a medical screening form reviewed and approved by an aerospace physician, based on a cardiologist’s assessment of their exercise electrocardiogram. All experimental procedures complied with the principles outlined in the Declaration of Helsinki, and were approved by the French national ethics committee *Comité de Protection des Personnes Île-de-France II* (No. 2023-A02145-40).

#### Parabolic flight exposure

Parabolic flights were conducted during the VP177 and VP181 campaigns organized by the *Centre National d’Études Spatiales* (CNES, France) aboard the Airbus A310 Zero G, operated by Novespace in Mérignac. Each flight lasted 2.5 to 3 hours and comprised 31 parabolic maneuvers, consisting of successive gravity phases: hypergravity (1.8 g), microgravity (0 g), and a return to hypergravity (1.8 g), interspersed with periods of normogravity during steady flight. Each parabola lasted approximately one minute, followed by a two-minute interval between consecutive parabolas. Five-minute breaks were scheduled after every five parabolas, and an extended eight-minute break occurred midway through the session. During the flight, participants performed two standard vestibulo-motor tasks distributed throughout the parabolic session. These tasks included a *vestibulo-ocular reflex* experiment and a *vestibular evoked myogenic potential* experiment, which are not reported here (see Bonnard et al., 2025 for further details). Conducted independently of the parabolic flights, these tasks are not known to induce motion sickness.

Although scopolamine is commonly administered before parabolic flights to mitigate motion sickness by dampening vestibular integration (Bestaven, 2016 ; Weerts, 2015), no medication was provided in the present study.

#### Virtual reality exposure

Participants stood upright on a platform and wore a virtual reality headset for the sensory conflict procedure (*MotionVR*, Virtualis, Montpellier, France). They were exposed to two different simulations: (1) a front passenger view of a car driving along a winding mountain road, and (2) a tunnel environment rotating randomly in orientation every 10 s and alternating between forward and backward motion every 8 s. Each simulation lasted 5 min and was repeated twice.

A continuous dialogue was maintained between the participant and the experimenter to monitor subjective sensations throughout the exposure. The procedure was terminated when participants reported moderate nausea or discomfort, corresponding to a score of 4 on the Motion Sickness Severity Scale (Czeiler, 2023), a validated questionnaire designed to assess real-time motion sickness symptoms. Those who did not experience symptoms remained in the simulation for a total of 20 min.

#### Objective assessment of MS: Physiological measures

In the VR experiment, body surface temperature, skin color, salivary cortisol, and blood pressure were measured at three time points: before the start of the pre-tests, immediately after cessation of VR exposure, and after completion of the post-tests (Figure 1B, upper panel). The pre- and post-tests consisted of four different sensory assessments: Vestibular Evoked Myogenic Potential, Vestibulo-Ocular Reflex, Sensory-Organisation Test and Optokinetic Nystagmus. These tests are not known to be nauseogenic (for more details, see Bonnard et al., 2025). Each pre- and post-test session lasted approximately 30 minutes, whereas the VR exposure lasted a maximum of 20 minutes. In the PF experiment, body surface temperature, skin color, and blood pressure were assessed before the parabolic session, during the first, mid-flight, and last breaks, and after the parabolic session (Figure 1B, lower panel). Salivary cortisol was collected only before the parabolic session, at mid-flight, and immediately after the final parabola. Heart rate and skin conductance were continuously recorded throughout both sessions.

**Figure 1.**
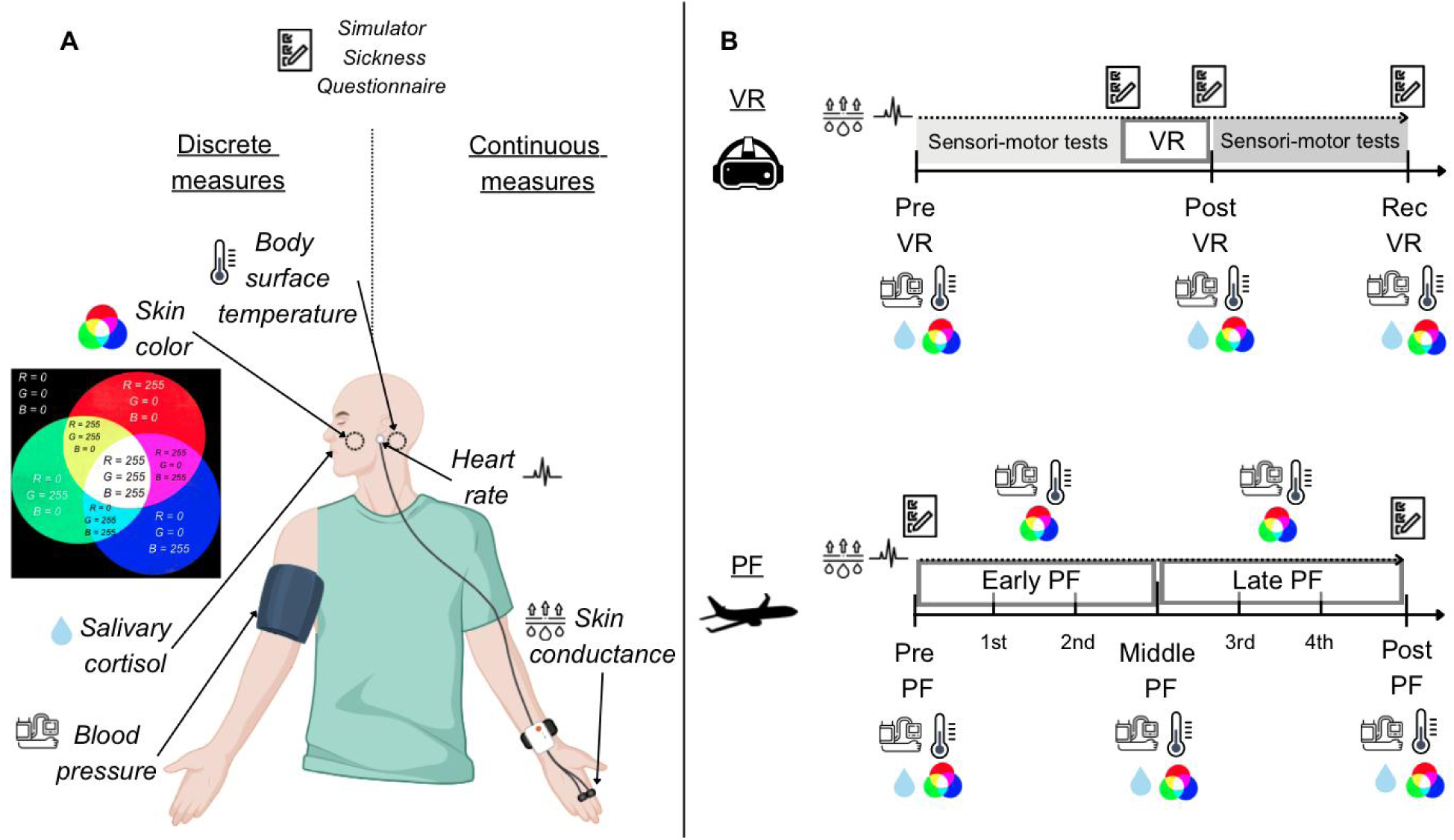
Description of objective physiological variables (A) and protocol schedules (B). Pre refers to the measurements taken at the start of the protocol, before any pre-tests or parabolas. Post refers to the measurements taken following exposure to the sensory conflict paradigm (either VR or PF). Middle refers to the measurements collected midway through the flight in PF, after the completion of 16 parabolas, whereas Rec refers to the recovery period at the end of the protocol when final measurements were taken. VR: virtual reality; PF: parabolic flight.

Body surface temperature was measured at the mastoid process behind the right ear using an infrared thermometer (DT8806H, HT Instruments, Faenza, Italia) (Figure 1A).

Skin color was assessed at the center of the left cheek using a colorimeter (ColorReader EZ, DC10-3, Datacolor GmbH, Marl, Germany). This device captures color by placing the sensor directly on the skin surface and illuminating it with an integrated LED, thereby avoiding interference from ambient light. It was very small, easy to handle and straightforward to perform with minimized potential noise from ambient lighting. The RGB color scale was used for analysis, quantifying the intensity of the red, green, and blue channels, where 0 indicates no intensity (dark) and 255 indicates maximum intensity (light).

Salivary cortisol was collected through passive drool into 1 mL Eppendorf tubes, until a volume of 0.5 mL was obtained (Figure 1A). Samples were immediately frozen at – 20 °C after the completion of the participant’s protocol. All samples were later analyzed in a single batch using a Cortisol ELISA test (Salivary Cortisol Enzyme Immunoassay Kit, Salimetrics, Carlsbad, USA). Results from the enzymatic reaction was read with the multimode plate reader Clariostar Plus (BMG Labtech, Champigny s/Marne, France).

Blood pressure, including systolic and diastolic values, was measured using a tensiometer (HEM-7154-E, Omron, Kyoto, Japan). Measurements were taken on the right arm while participants were seated, during both the VR and PF paradigms (Figure 1A).

Skin conductance was continuously monitored at 15.9 Hz using the Shimmer3 GSR+ device (Shimmer, Dublin, Ireland), capable of detecting signals ranging from 8 kΩ to 4.7 MΩ (125 µS to 0.2 µS). Two electrodes were placed on the proximal phalanges of the index and middle fingers of the left hand to record galvanic skin response (GSR) (Figure 1A). Skin conductance was automatically computed by the device as the inverse of skin resistance and expressed in Siemens (S).

Heart rate was derived from a photoplethysmography (PPG) signal recorded by the Shimmer3 GSR+ device using an optical pulse sensor, placed on the participant’s left earlobe (Figure 1A). From PPG signal, we also extracted the interbeat intervals (IBIs) and heart rate variability (HRV), defined as the standard deviation of the IBIs.

Skin conductance and heart rate were analyzed in MATLAB R2024b (The MathWorks, Natick, USA). Raw skin conductance signals were low-pass filtered using a fourth-order Butterworth filter with a 0.1 Hz cutoff frequency to remove noise related to finger movements. Heart rate data were cleaned by excluding extreme values, specifically those below 30 bpm or above 230 bpm. Representative examples of the filtered data are shown in Figure 2.

**Figure 2.**
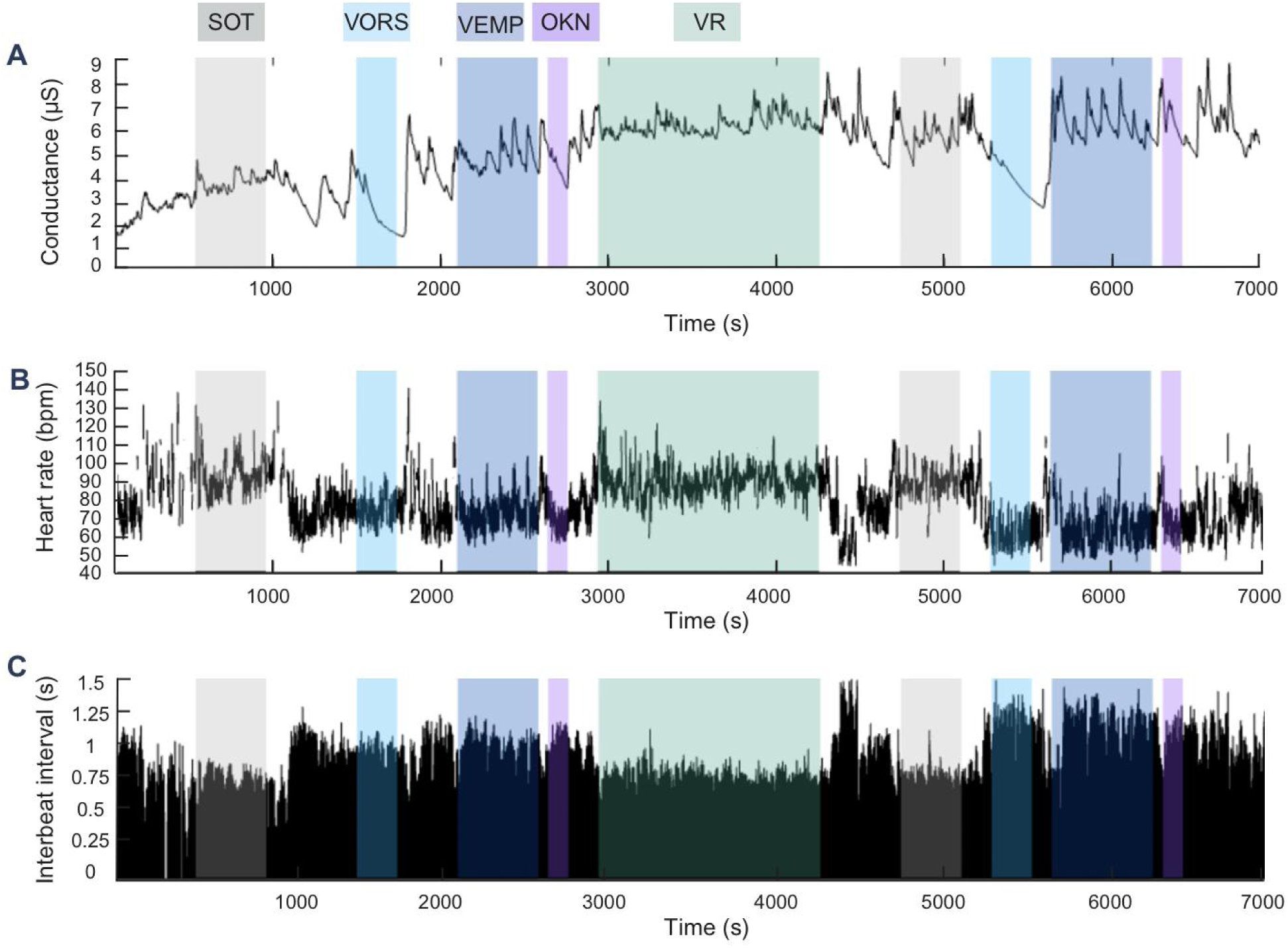
Example of continuous recording of skin conductance (A), heart rate (B) and interbeat-interval (C), from the VR experiment. Representations correspond to filtered data. Overlapping translucent color boxes represent the time portion were a sensory test have been performed. SOT, VOR, VEMP and OKN were performed both during pre and post-tests. SOT: sensory organization test, VOR: vestibulo-ocular reflex, VEMP: vestibulo-evoked myogenic potential, OKN: optokinetic nystagmus.

#### Subjective assessment of MS: questionnaires

The Motion Sickness Susceptibility Questionnaire (MSSQ; Golding, 2006) was completed before participants performing any paradigm. This questionnaire includes nine transport situations, each rated on a severity scale ranging from “Never sick” to “Frequently felt sick,” to assess how often an individual experiences motion sickness in such contexts— both during childhood (before 12 years old, MSA subscore) and adulthood (within the past 10 years, MSB subscore). These two subscores are then combined to yield a global susceptibility score. The MSSQ provides an estimate of how susceptible an individual may be to challenging motion environments.

The Simulator Sickness Questionnaire (SSQ, Kennedy, 1993) was completed before and after the virtual sensory conflict procedure, and before and after the completion of the parabolic session (Figure 1B). This questionnaire comprises 16 symptom descriptions, each accompanied by a severity scale ranging from ’not at all’ to ’severe,’ to assess the intensity of experienced symptoms. These symptoms can subsequently be categorized into two distinct groups: those related to gastrointestinal discomfort and nausea, and those associated with central and oculomotor functions. SSQ relevance lies in the difference between the baseline and post-exposure scores, as the phrasing prompts participants to report how they feel at the moment they complete the form.

The Motion Sickness Assessment Questionnaire (MSAQ) was also administered at the end of the parabolic flight session to better characterize symptom types. The MSAQ is a 16-item questionnaire, with responses scored from 1 (“not at all”) to 9 (“severely”), and provides subscale scores for four symptom domains: gastrointestinal, sopite, peripheral, and central. This questionnaire encompasses a wider range of symptoms and is not limited to a specific context. Its phrasing instructs participants to report how they felt during the exposure rather than at the moment of completing the form, meaning that the relevance of the MSAQ score relies on a single administration conducted after exposure.

#### Statistical analysis

Participants were objectively classified in two subgroup, *sick* and *non-sick*, based on the attainment of two predefined critical thresholds. In the VR paradigm, this threshold corresponded to the premature termination of the session—participants reported being unable to continue without imminently vomiting, typically within one minute. In the PF paradigm, participants were categorized as motion sick if they actually vomited, representing a physiological endpoint indicative of a maximal motion sickness response.

To compare physiological states across exposure durations and between sick and non-sick groups, we conducted two-way repeated-measures ANOVAs, using time points as the within-subjects factor and sickness as the between-subjects factor. Due to differences in time points and sickness profiles between the VR and PF paradigms, separate ANOVAs with Bonferroni correction were performed for each dataset. Prior to running the ANOVAs, homogeneity of variance and sphericity were verified and satisfied in all analyses (p > 0.05). Effect sizes for ANOVA results were reported as partial eta squared (η²), calculated as the ratio of the effect sum of squares to the sum of the effect and error sum of squares.

Independent *t*-tests were used to examine differences in subjective motion-sickness questionnaire scores in VR and PF. The grouping variable was sickness status, and the dependent variables were the Post–Pre differences in SSQ and MSAQ total scores and subscores. Normality was assessed using the Shapiro–Wilk test and homogeneity of variance using Levene’s test. When normality was not met in either group, a non- parametric Mann–Whitney *U* test was used instead of the Student *t*-test.

To determine whether physiological changes were similar across the two paradigms for each participant, we correlated the post–pre differences in physiological markers obtained in the VR condition with those from the PF condition (Spearman’s ρ). We also examined whether the physiological parameters measured before (Pre) exposure to either VR or PF were associated with motion sickness questionnaire scores (SSQ, MSAQ).

To evaluate the predictive relationships between baseline physiological measures and changes in MS symptoms, we conducted linear regression analyses. Specifically, we examined the association between baseline skin temperature, skin conductance, salivary cortisol concentration, and changes in SSQ scores following exposure. All assumptions (linearity, independence, homoscedasticity, and normality of residuals) were checked and met.

All statistical analyses were conducted using JASP (Version 0.95.4). Results are presented as means ± standard errors (SE) unless specified within the text. Boxplots in figures represent medians ± standard errors (SE).

## RESULTS

### Evolution of perceived MS following exposure to VR and PF

In the VR condition, 38% of participants (8 out of 21) stopped the simulation due to severe motion sickness. In contrast, 57% of participants (12 out of 21) vomited and were considered sick during the PF condition. Overall, 29% of participants (6 out of 21) experienced sickness in both paradigms, while 33% (7 out of 21) remained non-sick across both conditions.

In VR, the difference in total SSQ scores (Post–Pre) was significantly greater in sick participants compared to non-sick ones (Sick = 11.13 ± 1.11, Non-sick = 1.46 ± 1.37; U(19) = 6.00, p < 0.001). Among SSQ subscores, only the nausea subscore showed a similar pattern, with sick participants reporting higher gastrointestinal symptoms (Post–Pre Nausea SSQ: Sick = 10.13 ± 1.08, Non-sick = 1.39 ± 0.94; t(19) = 5.97, p < 0.001; Figure 3A), whereas no significant difference was found for oculomotor symptoms (U(19) = 72.5, p = 0.129).

**Figure 3.**
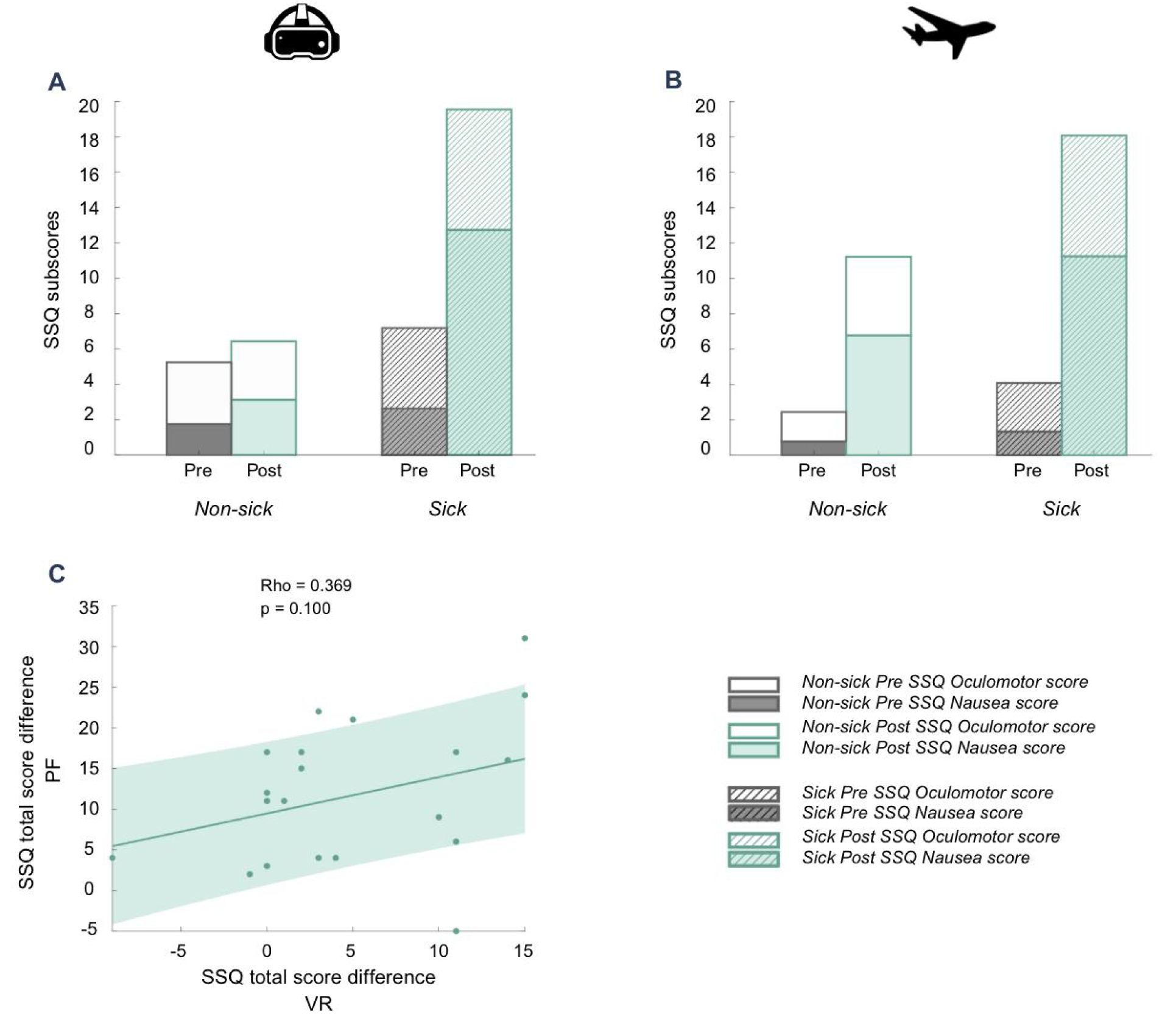
Subjective motion sickness before and after exposure to VR (A) and PF (B). For each bar, the Nausea subscore is plotted at the base and the Oculomotor subscore is superimposed above it. In panel (C), the correlation between subjective consequences across both paradigms was assessed using the post-minus-pre Total SSQ scores for each participant who took part in both the VR and PF protocols. Spearman correlations ; *p < 0.05, **p < 0.01. PF: parabolic flight, VR: virtual reality, SSQ: Simulator sickness questionnaire.

In the PF condition, the difference in total SSQ scores between sick and non-sick participants was not significant (Sick = 14.00 ± 2.46, Non-sick = 8.78 ± 2.87; t(19) = –1.39, p = 0.182; Figure 3B). However, post-flight MSAQ total scores were significantly higher in sick participants compared to non-sick ones (Sick = 59.55 ± 4.19, Non-sick = 33.64 ± 4.40; t(19) = –4.201, p < 0.001). Significant differences were also observed in the Gastrointestinal (Sick = 10.77 ± 0.67, Non-sick = 5.36 ± 1.00; U(19) = –4.66, p < 0.001), Central (Sick = 11.14 ± 0.99, Non-sick = 6.50 ± 1.12; t(19) = –3.09, p = 0.006), and Sopite (Sick = 7.23 ± 0.68, Non-sick = 3.36 ± 0.51; t(19) = –4.28, p < 0.001) subscores, with sick participants consistently reporting higher symptom levels.

The differences in SSQ scores between the VR and PF conditions showed a trend toward a positive correlation (ρ = 0.369, p = 0.100, n = 21; Figure 3C). This finding aligns with our previous results, which indicated that fewer than one-third of participants experienced sickness in only one condition, while two-thirds responded similarly in both contexts.

### Evolution of physiological parameters during exposure to VR and PF

Salivary cortisol: The increase in salivary cortisol was pronounced after PF exposure (Pre-PF = 0.30 ± 0.06, Post-PF = 1.84 ± 0.32 ; F(1,18) = 38.696, p < 0.001, η² = 0.683 ; Figure 4A), and this evolution did not significantly differ between sick and non-sick individuals in PF (F(1,18) = 1.937, p = 0.181). In VR, the Time points x Sickness interaction was significant (F(1,18) = 4.400, p = 0.050, η² = 0.196), and post-hoc analysis revealed a tendancy towards an increase in cortisol level for sick participants only (Sick Pre = 0.46 ± 0.10, Sick Post = 0.86 ± 0.21; t(7) = -2.74, p = 0.082), whereas this level remain stable for Non-Sick(Non-Sick Pre = 0.32 ± 0.47, Non-Sick Post = 0.33 ± 0.05 ; t(12) = -0.034, p = 1). The change in cortisol levels (Post-Pre) in the PF condition did not significantly correlate with the corresponding change in VR (ρ = 0.405, p = 0.086, n = 19; Figure 4B). This suggests that participants who exhibited stronger physiological stress responses in one condition were not necessarily the same individuals who did so in the other condition.

**Figure 4.**
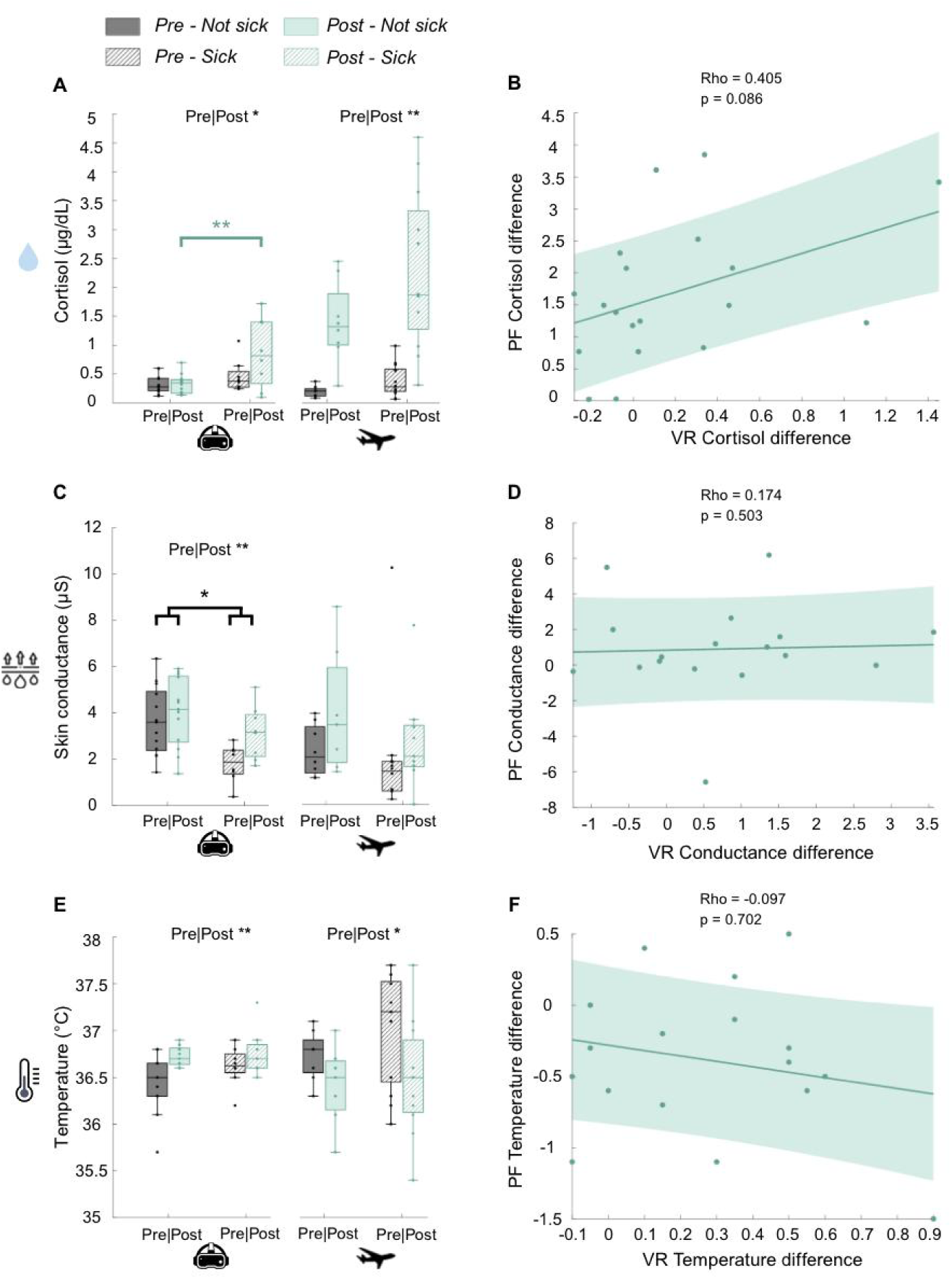
Evolution of salivary cortisol (A), skin conductance (C) and temperature (E) following exposure in VR and PF. Pre-values are presented in dark gray and post-values in green. Sick participants values are represented in striped boxes whereas non-sick individuals are represented in full boxes. Pre-values for physiological parameters in both paradigms correspond to measurements taken before the experiment. Post-values for salivary cortisol in VR were taken after the recovery period (accounting for cortisol peak latency ∼ 30 min), whereas post-PF values were collected after completion of all parabolas (parabolic session total duration ∼ 1h30). For skin conductance and temperature, post-values correspond to measurements immediately after VR or PF exposure. For correlations, differences were calculated as post-values minus pre-values (B, D, F). Two-way repeated measures ANOVA with Bonferroni post-hoc tests ; Spearman correlations ; *p < 0.05, **p < 0.01. PF: parabolic flight, VR: virtual reality.

Skin conductance: A significant effect of Time point was observed in VR (F(1, 19) = 13.43, p = 0.002, η² = 0.414), with increased values post-exposure compared to pre- exposure (Pre-VR = 2.73 ± 0.35; Post-VR = 3.60 ± 0.41 ; Figure 4C). A significant main effect of Sickness group was also found (F(1, 19) = 6.21, p = 0.022, η² = 0.246), with lower conductance values in sick individuals compared to non-sick participants (Sick = 2.48 ± 0.34 ; Non-sick = 3.85 ± 0.41). The Time points x Sickness interaction was not significant in VR (F(1, 19) = 4.032, p = 0.059) but showed a tendency towards greater skin conductance increase post-exposure in sick participants. In PF, neither Time points (F(1,15) = 2.144, p = 0.164) nor Sickness (F(1,15) = 0.921, p = 0.353) presented significant differences for skin conductance, and there was no interactions between these factors (F(1, 15) = 0.528, p = 0.479). Similarly to cortisol, changes in skin conductance elicited by the PF and the VR conditions were not significantly correlated (ρ = 0.174, p = 0.503, n = 17 ; Figure 4D).

Temperature: Surface skin temperature was significantly increased after VR exposure (Pre-VR = 36.54 ± 0.08, Post-VR = 36.75 ± 0.06 ; F(1, 19) = 9.974, p = 0.005, η² = 0.344; Figure 4E), whereas it decreased following PF (Pre-PF = 36.81 ± 0.14, Post-PF = 36.45 ± 0.18 ; F(1, 16) = 7.779, p = 0.013, η² = 0.327 ; Figure 4E). Neither Sickness nor interaction between Time points and Sickness were observed in both paradigms (all p>0.05). As with previous variables, intra-individual changes in skin temperature were not correlated between the two types of exposure (ρ = –0.097, p = 0.702, n = 17; Figure 4F).

Blood pressure: In the VR condition, sick participants exhibited significantly lower systolic pressure compared to non-sick participants (Sick = 108.19 ± 3.03, Non-Sick = 121.06 ± 3.03 ; t(19) = 2.989, p = 0.008, η² = 0.320 ; Figure 5A). This difference was independent of VR exposure effects, as no significant main effect of Time Points (Pre-VR = 115.04 ± 4.04, Post-VR = 114.21 ± 2.35, p = 0.761) or Time Points × Sickness interaction (F(1,19) = 0.201, p = 0.659) was observed. In contrast, systolic pressure significantly decreased after exposure in the PF condition (Pre-PF = 127.42 ± 5.28, Post-PF = 116.37 ± 4.34 ; F(1,10) = 8.787, p = 0.016, η² = 0.494 ; Figure 5A). However, there were no significant effect of Sickness (F(1,10) = 0.642, p = 0.642) a significant Sickness × Time Points interaction (F(1,10) = 0.096, p = 0.764). Individual differences in systolic blood pressure variation across the two protocols did not correlate (ρ = 0.151, p = 0.658, n = 11 ; Figure 5B).

**Figure 5.**
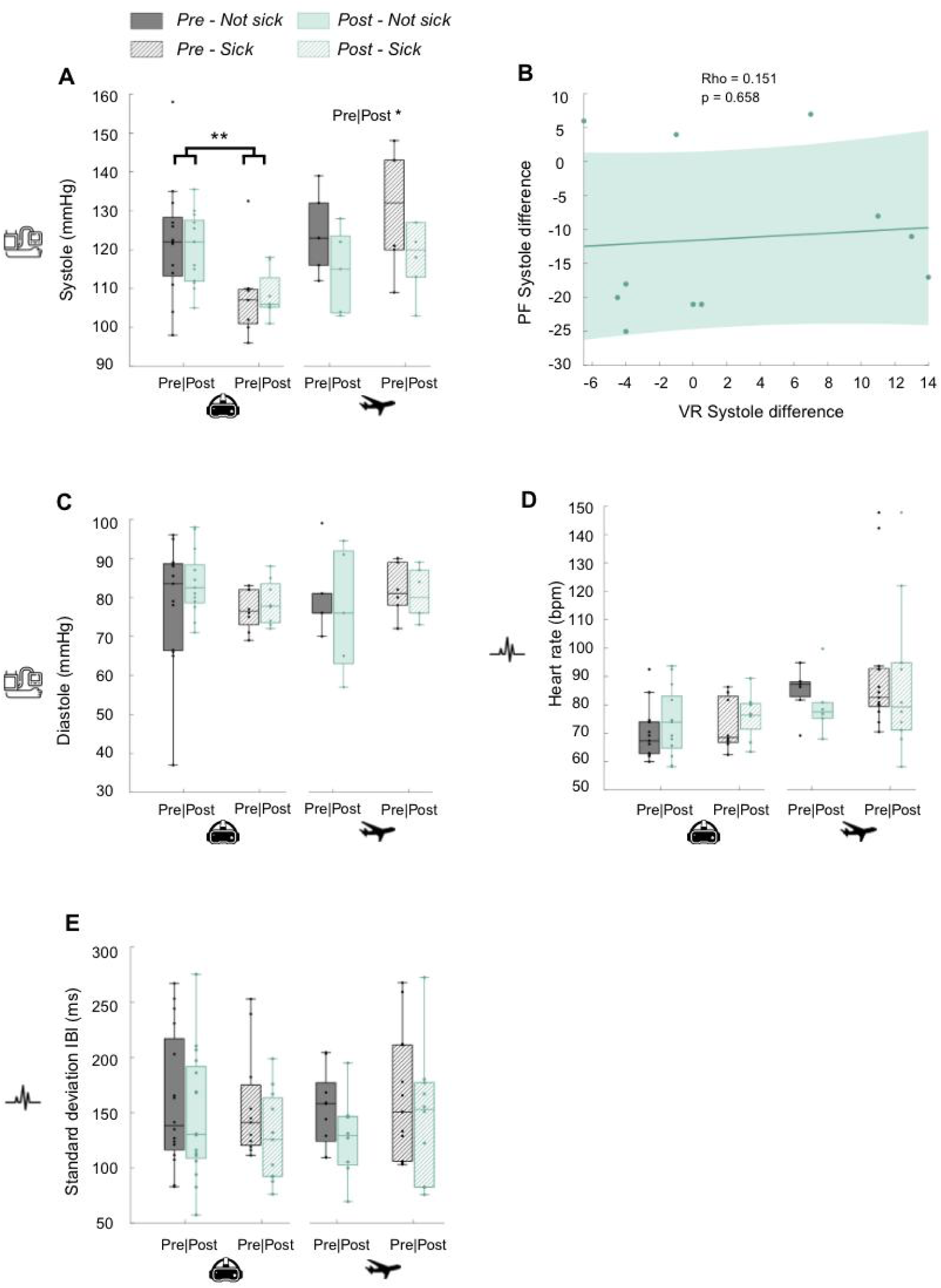
Evolution of systolic pressure (A), diastolic pressure (C), heart rate (D) and heart rate variability (E) following exposure in VR and PF. Pre-values are presented in dark gray and post-values in green. Sick participants values are represented in striped boxes whereas non-sick individuals are represented in full boxes. Pre-values correspond to measurements taken before the experiment. Post-values were taken right after exposure to VR or parabolic session. Differences were calculated as post-values minus pre-values for correlation (B). Two-way repeated measures ANOVA with Bonferroni post-hoc tests ; Spearman correlations ; *p < 0.05, **p < 0.01. PF: parabolic flight, VR: virtual reality, IBI: interbeat interval.

For diastolic pressure (Figure 5C), no significant effects of Time Points, Sickness, or their interaction were observed in either paradigm (all p > 0.05 ; Pre-PF = 80.75 ± 3.45, Post-PF = 78.77 ± 4.98 ; PF-Sick = 81.33 ± 2.76, PF-Non-Sick = 84.88 ± 9.37 ; Pre-VR = 77.55 ± 3.17, Post-VR = 81.31 ± 2.19 ; VR-Sick = 77.78 ± 1.96, VR-Non-Sick = 81.08 ± 3.41).

Heart rate: No significant effects of Time Points, Sickness, or their interaction were observed were observed in both VR (all p > 0.05 ; Pre-VR = 71.72 ± 2.95, Post-VR = 74.99 ± 3.09) and PF (all p > 0.05 ; Pre-PF = 89.39 ± 5.73, Post-PF = 84.24 ± 6.50 ; Figure 5D).

Similarly, heart rate variability (HRV) displayed no significant effects of Time Points, Sickness, or their interaction in both protocols (all p > 0.05 ; Pre-PF = 149.09 ± 14.28, Post-PF = 138.51 ± 17.13 ; PF-Sick = 147.09 ± 18.10, PF-Non-Sick = 140.51 ± 13.30 ; Pre-VR = 158.03 ± 15.02, Post-VR = 137.40 ± 13.41 ; VR-Sick = 141.80 ± 13.60, VR-Non- Sick = 153.63 ± 14.82; Figure 5E).

Skin color: In the VR paradigm, sick participants exhibited a reddish facial coloration at the end of the VR protocol. The red channel showed a significant Time points × Sickness interaction (F(2, 50) = 5.537, p = 0.007, η² = 0.181) with red values significantly lower after the recovery period compared to immediately post-exposure in sick participants (Post-VR = 100.93 ± 0.27, Rec-VR = 98.83 ± 0.42, t(15) = 4.014, p = 0.003), whereas non-sick participants did not exhibit a significant change (Figure 6A).

**Figure 6.**
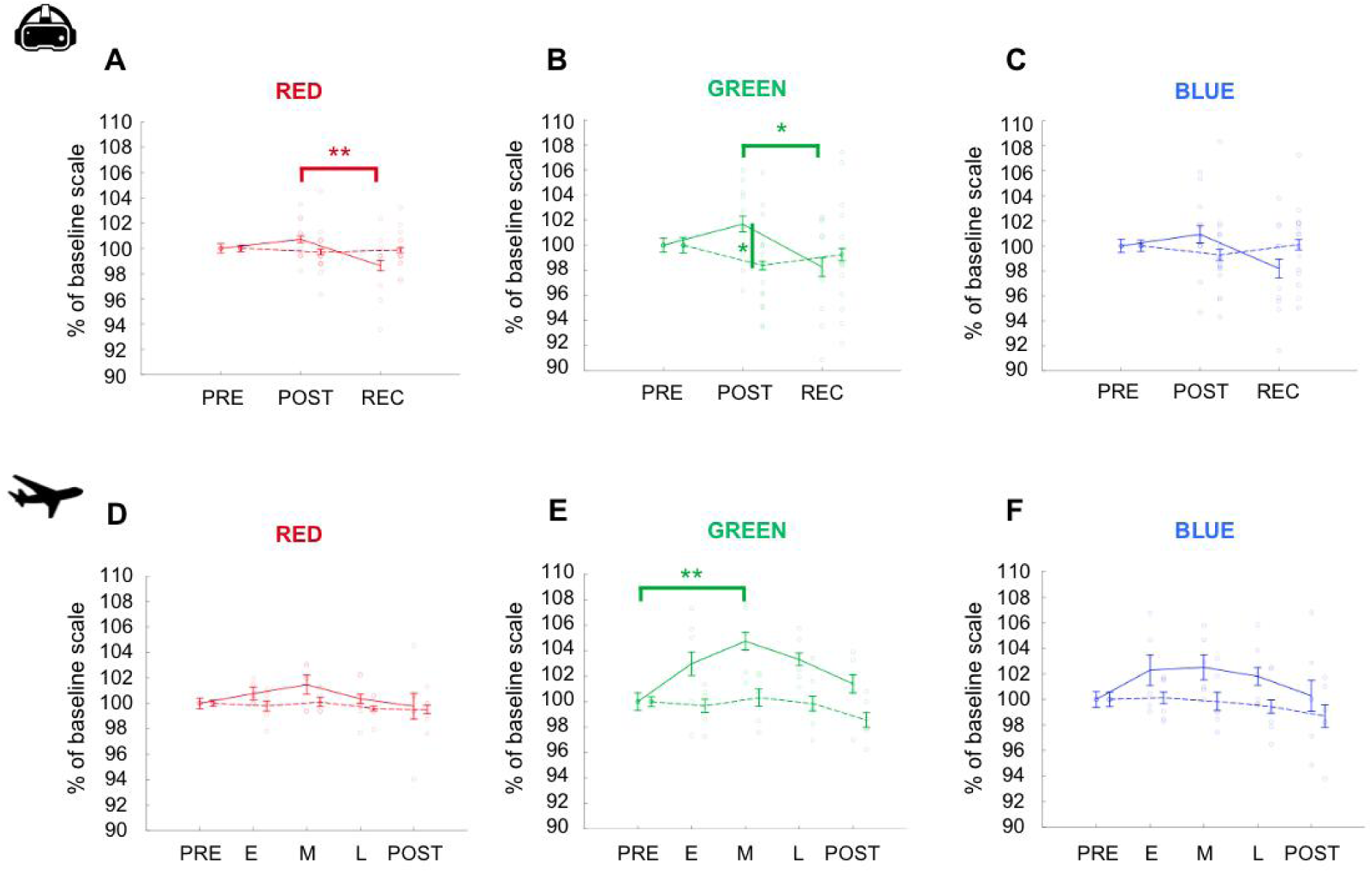
Skin RGB scale evolution over time in VR (A-C) and in PF (D-F). Sick participants are represented in full line while non-sick participants are represented in dotted line. Each color channel of the RGB scale is calculated as a percentage of the baseline, corresponding of the color channel value before experiment. Two-way repeated measures ANOVA, Bonferroni post-hoc test. *: p<0.05 ; **: p<0.01. PF: parabolic flight, VR: virtual reality, RGB: Red-Green-Blue scale, PRE (in VR): before experiment, PRE (in PF): before parabolas, POST (in VR): right after VR exposure, POST (in PF): right after parabolic session, REC: recovery phase at the end of protocol, E: early flight, M: middle flight, L: late flight.

The green channel showed a significant Time points×Sickness interaction (F(2, 50) = 5.228, p = 0.009, η² = 0.173). Sick participants exhibited increased facial greenness after the VR exposure, with higher green values compared to non-sick individuals in post-VR only (Sick Post-VR = 101.698 ± 0.63, Non-Sick Post-VR = 99.15 ± 0.35 ; t(25) = -3.161, p = 0.04 ; Figure 6B). No Pre/Post effect was found for this channel (Pre = 100.45 ± 0.59, Post = 100.42 ± 0.49, t(25) = 0.034, p = 1).

The blue channel also showed a significant Time points × Sickness interaction (F(2, 50) = 3.658, p = 0.033, η² = 0.128) but post-hoc comparisons were not significant (p > 0.05 ; Figure 6C).

In the PF paradigm, neither the red nor the blue channels showed significant effects of Time Points or Sickness (all p > 0.05 ; Figure 6D&F). In contrast, the green channel showed a significant Time points × Sickness interaction (F(4, 40) = 2.633, p = 0.048, η² = 0.208). Green values were significantly increased from pre-parabolas to mid-flight in sick participants (Sick: Pre-PF = 97.60 ± 0.67, Middle-PF = 102.20 ± 0.67 ; t(10) = -4.594, p = 0.002). This effect was not observed in non-sick participants (Non-Sick: Pre-PF = 100.33 ± 0.39, Middle-PF = 100.63 ± 0.68 ; t(10) = -0.299, p = 1) (Figure 6E). The decrease from mid-flight to post-parabolas was not significant in either sick or non-sick group (all p > 0.05).

Similarly to other physiological parameters, none of the variations in skin color channels in one paradigm was correlated to changes in color channels in the other (all p>0.05).

### Relation between identified physiological markers and perceived MS

Based on the observed significant physiological changes following exposure to both VR and PF, we hypothesized that these specific variations might reflect reported subjective MS severity and symptomatology.

Cortisol: In VR, changes in salivary cortisol were significantly correlated with changes in Total SSQ scores (ρ = 0.573, p = 0.008, n = 20) and Nausea subscores (ρ = 0.643, p = 0.002, n = 20 ; Figure 7A), indicating that greater symptom severity was associated with higher increase of cortisol. In PF, cortisol changes correlated only with the MSAQ Gastrointestinal subscore (ρ = 0.475, p = 0.034, n = 20 ; Figure 7B), with participants reporting stronger GI symptoms (e.g. nausea) exhibiting higher increase of cortisol.

**Figure 7.**
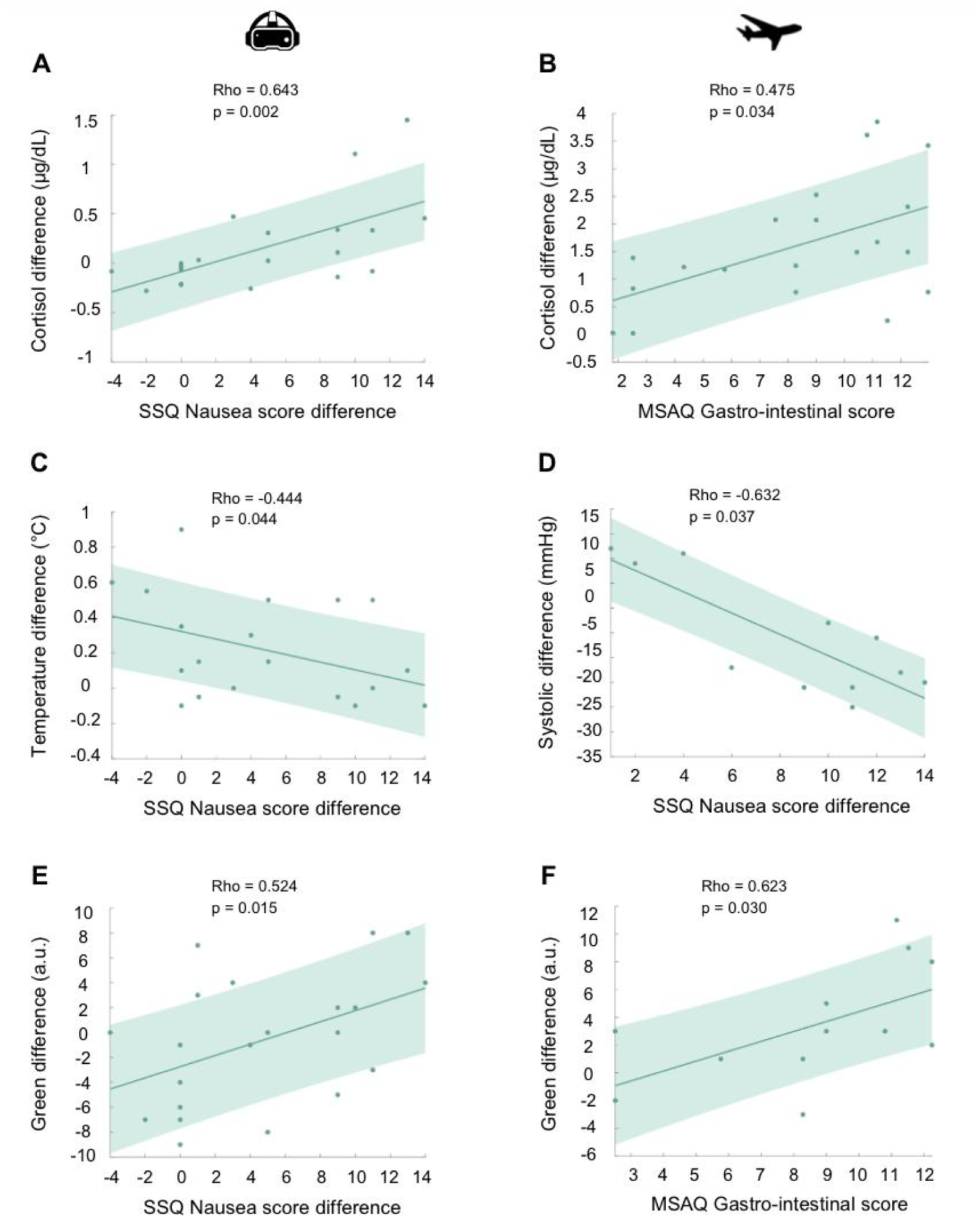
Relationship between perceived MS and physiological evolution in VR (A, C, E) and PF (B, D, F). In the PF condition, differences in cortisol levels and systolic blood pressure were calculated as the change between post-parabolic and pre-parabolic measurements. For the in-flight green-channel signal, differences were computed using mid-flight values relative to pre-parabolic baseline values. In the VR condition, cortisol differences corresponded to recovery values minus pre-protocol values, whereas temperature and green- channel differences were calculated as pre-protocol values minus post-VR values. Spearman correlations. MSAQ: motion sickness assessment questionnaire, SSQ: simulator sickness questionnaire.

Skin conductance: In VR, changes in skin conductance were not correlated with any SSQ score variations (all p > 0.05).

Skin temperature: In VR, smaller skin surface temperature variations were associated with increased nausea (SSQ Nausea: ρ = –0.444, p = 0.044, n = 21; Figure 7C). In PF, no correlations were found between skin temperature changes and questionnaire scores (all p > 0.05).

Systolic blood pressure: In VR, systolic pressure changes were unrelated to SSQ total or subscore differences (all p > 0.05). In PF, reductions in systolic pressure were correlated with increased nausea (SSQ Nausea: ρ = –0.632, p = 0.037, n = 11 ; Figure 7D) and showed a strong trend with higher MSAQ total scores (ρ = –0.592, p = 0.055, n = 11). Participants who experienced greater nausea-like symptoms displayed larger systolic reductions post-flight.

Heart rate variability: HRV changes were not significantly correlated with questionnaire scores in VR and in PF (all p > 0.05).

Green RGB channel: Increases in the green channel were consistently associated with higher nausea symptoms in both VR and PF. In VR, green-channel differences correlated with changes in Total SSQ (ρ = 0.470, p = 0.031, n = 21) and Nausea scores (ρ = 0.527, p = 0.015, n = 21; Figure 7E). In PF, pre–to–mid-flight green-channel changes correlated with MSAQ Gastrointestinal symptoms (ρ = 0.623, p = 0.030, n = 12 ; Figure 7F) and showed a strong trend with Sopite symptoms (ρ = 0.559, p = 0.059, n = 12).

During PF, we recorded the parabola number at which sick participants first vomited and assigned parabola 30 to participants who did not become sick. The parabola of first vomit was significantly correlated with in-flight MSAQ general (ρ = –0.636, p = 0.002, n = 21), Gastrointestinal (ρ = –0.591, p = 0.005, n = 21), Central (ρ = –0.566, p = 0.007, n = 21), and Sopite symptoms (ρ = –0.817, p < 0.001, n = 21). Participants who became sick earlier in the flight reported stronger motion-sickness symptoms. In contrast, with physiological variations, the parabola of first vomit did correlate only with green channel change from pre-parabola to mid-flight (ρ = –0.819, p = 0.001, n = 12). The sooner the participant vomited during the first part of the flight, the greener he was mid-flight.

Although we also recorded the time at which participants stopped the VR simulation, too many individuals completed the full 20-minute session, violating the assumptions required for correlation analyses. As a result, correlations with VR exposure duration were not computed.

### Predictors of Motion Sickness

Regarding subjective susceptibility based on past experiences, MSSQ total scores and subscores (MSA and MSB) were not correlated with any physiological changes in either paradigm (all p > 0.05). In addition, MSSQ total scores did not correlate with changes in Total SSQ scores in VR or PF, nor with Total MSAQ scores in PF.

However, adulthood MSSQ scores were positively correlated with changes in Oculomotor SSQ symptoms in VR (ρ = 0.465, p = 0.034, n = 21) and with Central MSAQ symptoms in PF (ρ = 0.457, p = 0.037, n = 21). None of these associations remained significant when tested in linear regression models assessing predictive value.

In the VR paradigm, only the baseline salivary cortisol level was positively correlated with the increase in SSQ oculomotor symptoms following exposure (ρ = 0.503, p = 0.020, n = 21 ; Table 1 line 1). Higher cortisol level before protocol was associated with higher oculomotor symptoms post-exposure. However, this elevated cortisol level did not predict the variance in oculomotor scores due to VR exposure (β = 0.210, SE = 1.957, R^2^ = 0.044, p = 0.362).

**Table 1.**
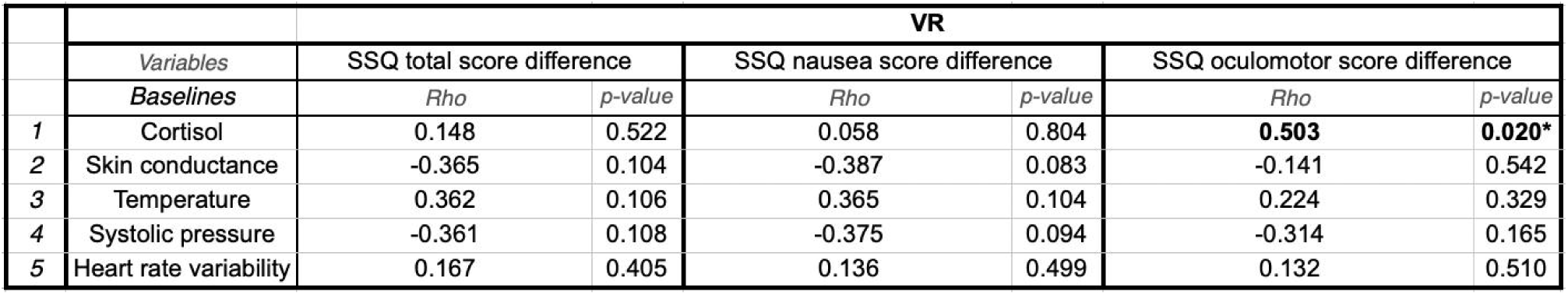
Correlation matrix between baseline physiological parameters and subjective motion-sickness scores in VR. Baseline measures were taken before starting the VR protocol. Spearman correlations. *: p<0.05 ; **: p<0.01. SSQ = Simulator sickness questionnaire.

In the PF condition, among baseline physiological parameters, temperature, skin conductance and heart rate variability showed an interesting correlation with perceived sickness post-flight. MSAQ Central score was significantly correlated with pre-parabola surface skin temperature (ρ = 0.59, p = 0.005, n = 21; Table 2 line 3). Greater pre-parabola temperature was linked to higher central symptoms post-flight. Specifically, pre-parabola temperature explained 31.5% of the variance in central symptoms (β = 0.561, SE = 1.542, *R*² = 0.315, *p* = 0.008). Pre-parabola skin conductance was also positively correlated to peripheral post-flight MSAQ subscore (ρ = 0.472, p = 0.031, n = 21; Table 2 line 2) but did not predicted the pre-to-post variance of those symptoms (β = 0.236, SE = 0.177, R ^2^ = 0.056, p = 0.304). Similarly, pre-parabola heart rate variability was positively correlated to peripheral post-flight MSAQ subscore (ρ = 0.473, p = 0.030, n = 21; Table 2 line 5) and did predict 22.2% of the variance in those symptoms (β = 0.511, SE = 0.101, *R*² = 0.222, *p* = 0.018).

**Table 2.**
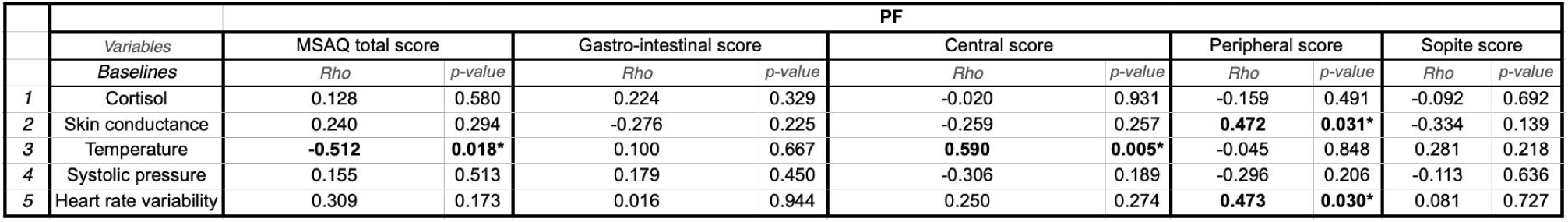
Correlation matrix between baseline physiological parameters and subjective motion-sickness scores in PF. Baseline measures were taken in flight before parabolic session. Spearman correlations. *: p<0.05 ; **: p<0.01. MSAQ: motion sickness assessment questionnaire.

## DISCUSSION

In this study, we investigated the evolution of physiological parameters across different motion-sickness–inducing paradigms, corresponding to virtual reality simulations and gravity alterations during parabolic flight. The purpose was to identify objective physiological markers of MS and to analyze the similarities and divergences of these markers between PF and VR paradigms. Additionally, this approach allowed for a comparative assessment of individual sensitivity to both paradigms, while also exploring potential objective predictors of MS.

Several physiological responses to motion sickness (MS) were consistent across both nauseogenic contexts, including elevated cortisol levels post-exposure, increased skin greenness, while heart rate, heart rate variability and diastolic blood pressure remained stable throughout the experiments. The post-exposure increase in cortisol in both paradigms and its consistent association with gastrointestinal symptoms were expected, given cortisol’s established role as a stress hormone linked to subjective motion sickness (Otto et al., 2006; Choukèr et al., 2010).

With regard to the differences between paradigms, we measured a decrease of systolic pressure after PF which was not observed in VR. This decrease in systolic blood pressure was strongly correlated with motion sickness (MS) intensity experienced during parabolic flight (PF). This result supports the hypothesis that changes in arterial pressure represent a specific symptom of vestibular-driven motion sickness. This aligns with previous observations of arterial blood pressure reduction following sinusoidal galvanic vestibular stimulation (Javaid et al., 2019) and centrifugation (Eiken et al., 2005).

We also observed trends in temperature that exhibited opposing patterns between the two paradigms. These results have to be interpreted cautiously, as both experimental setup and environmental factors could have influenced them. In VR, the headset was large and covered a substantial portion of the participants’ faces, leading to reports of feeling "hot". This resulted in a temperature increase across all participants, not only in those who developed cybersickness, contradicting previous literature that documented a decrease in skin temperature during cybersickness (Keshavarz et al., 2024; Arnold et al., 2019). In parabolic flight (PF), the aircraft cabin temperature fluctuated between 17 and 20°C, causing an approximate 3°C variation. Given that ambient temperature influences skin surface temperature (Liu et al., 2013; Mazdeyasna et al., 2023), these fluctuations likely affected participants’ skin temperature. Once again, although temperature is known to decrease with MS (Nobel, 2010; Nalivaiko et al., 2014), the temperature changes in PF occurred universally, regardless of sickness, suggesting a predominant influence of environmental factors. Taken together, these observations highlight the change in systolic blood pressure as the only relevant physiological marker distinguishing motion sickness induced by virtual reality from that induced by parabolic flight.

A key finding was the objective monitoring of skin color using colorimetry, which revealed an increase in skin greenness. To our knowledge, this is the first study to assess skin color changes across comparative motion sickness paradigms; previous research has relied primarily on subjective visual assessments. While only two studies have reported pallor increases using the L*a*b scale (Holmes, 2006; Li et al., 2022), our approach was sensitive enough to detect distinct changes in the red, green, and blue channels of the RGB scale—a more intuitive scale for interpreting skin color. Notably, the green channel was most affected by motion sickness, aligning with the colloquial description of "turning green" when ill. This increase in greenness followed prolonged sensory conflicts and correlated with the severity of gastrointestinal symptoms, particularly during parabolic flight where only sick participants exhibited increased green values mid-flight --the period which they vomitted. The green increase was modest in VR, likely because participants stopped the simulation before reaching the non-return point for vomiting.

Skin color results from the combined contributions of several chromophores: red (oxyhemoglobin), blue (deoxygenated hemoglobin), yellow–orange (carotene, an exogenous pigment), and brown (melanin) (Ortonne, 2012). While melanin and carotene are influenced by diet and light exposure, hemoglobin oxygenation can be directly modulated by physiological challenges. Several studies have shown that skin microcirculation increases during motion sickness, primarily to facilitate heat loss through thermoregulation (Nobel et al., 2012; Nalivaiko, 2018), and this has been monitored specifically in facial regions (Kolev et al., 1997). These studies also emphasize that vasoconstriction and vasodilation are largely region-specific and vary with the severity of motion sickness, with vomiting potentiating both pallor and vasoconstriction (Javaid et al., 2018). Given the substantial modulation of microcirculation during motion sickness, it is reasonable to hypothesize that skin blood oxygenation may decrease locally, resulting in a shift toward a blue–yellowish (i.e. greenish) hue. In this study, skin colorimetry—specifically the green channel—emerged as the most straightforward parameter to measure, and the physiological variable most closely aligned with subjective discomfort as evidenced by the strong and significant correlation between these measures across both paradigms.

Given the substantial sample size for a study involving parabolic flight and space-like motion sickness, it was particularly valuable to assess individual sensitivity to motion sickness using objective physiological measures. The objective was to determine whether individuals susceptible to motion sickness (MS) in one paradigm were similarly affected in the other, which could, for example, enable the prediction of space sickness susceptibility based on an individual’s history of terrestrial motion sickness. For all variables affected by either parabolic flight (PF) or virtual reality (VR) exposure—including cortisol levels, skin conductance, body temperature, systolic pressure, heart rate variability, and skin greenness—no significant correlation was observed between the effects in PF and those in VR. This suggests that individuals’ physiological changes to motion sickness (MS) in one paradigm are not necessarily similar in the other. This conclusion aligns with subjective assessments of discomfort, as the participants reporting discomfort differed between paradigms, evidenced by the lack of correlation between the Simulator Sickness Questionnaire (SSQ) scores in VR and post-flight Motion Sickness Assessment Questionnaire (MSAQ) scores. Behavioral data further support this finding, with no correlation between the first-vomit parabola in PF and the point of simulation termination in VR. Collectively, these physiological, behavioral, and subjective approaches align with prior research that, in small populations, failed to establish a relationship between ground- based assessment tests and space-flight susceptibility (Lackner and Dizio, 2006, for review).

Finally, we investigated the potential relationships between candidate predictors and perceived motion sickness in both protocols. The MSSQ global score failed to predict individual susceptibility to motion sickness in either paradigm, suggesting that past experiences of motion sickness do not reliably predict subsequent susceptibility in these contexts. This aligns with previous findings in parabolic flight studies (Golding et al., 2017) but contrasts with earlier VR research, where the MSSQ correlated with SSQ scores (Golding et al., 2021). Regarding objective predictors, baseline physiological values were analyzed prior to exposure. In VR, baseline cortisol was correlated with oculomotor symptoms, indicating that factors such as stress may influence centrally mediated symptoms. In PF, baseline skin conductance correlated with peripheral-type symptoms. However, neither baseline cortisol in VR nor baseline skin conductance in PF reliably predicted central and peripheral symptoms respectively. Only baseline skin temperature and baseline heart rate variability in PF were correlated with and predictive of variance in central and peripheral symptoms respectively. Higher baseline skin temperatures were associated with increased central symptoms post-flight, while the extent of temperature change showed no correlation with questionnaire scores. Notably, some participants exhibited skin surface temperatures around 38 °C, which is above the usual range and can induce thermal discomfort. This effect may stem from a disruption in circadian rhythms due to waking earlier than usual on the day of the flight, potentially impacting thermoregulation (Dijk et al., 2012). However, this hypothesis was not supported, as we did not find any correlation between baseline cortisol levels and baseline temperature in PF. Given that sleep quality is also known to influence motion sickness susceptibility (Kaplan et al., 2017; Altena et al., 2019; Bonnard et al., 2026b), further research is needed to clarify the interplay between sleep, circadian rhythm, vestibular integration, and motion sickness. Heart rate variability (HRV) exhibited a predictive value of just 22.2% for space motion sickness—a performance even lower than that of temperature. This finding indicates that HRV may not serve as a reliable predictor in the present dataset. Furthermore, the literature consistently highlights the need for caution when interpreting HRV metrics, given the persistent challenges in disentangling their association with sympathetic activity, parasympathetic activity, or the dynamic balance between the two (Malik, 1996). Consequently, the interpretation of this result remains both complex and of limited practical utility.

Interestingly, none of the baseline physiological parameters predicted the gastrointestinal symptoms, which were the predominant and most incapacitating symptoms, particularly in PF. One possible explanation is that many of the physiological parameters we measured may vary as a consequence of nausea rather than acting as factors influencing its emergence, unless these parameters fall outside their usual range. Nausea is primarily mediated through three major pathways: the area postrema in the brainstem (Borison & Wang, 1953; Miller & Leslie, 1994), vestibular afferents (Lackner, 2014; Oman & Cullen, 2014), and visceral afferents (Stern et al., 2011; Babic & Browning, 2015). The area postrema was likely not involved in our study, as no emetogenic substances were administered. Vestibular sensitivity, assessed in our study, did not predict motion sickness in PF, although it showed potential predictive value in VR (Bonnard et al., 2026a). The visceral pathway remains the least understood. Given the proposed presence of graviceptors in the viscera (Mittelstaedt, 1996), the stimulation of these receptors during parabolic flight (PF) may have contributed to the observed inter-individual differences in motion sickness susceptibility (Gierke & Parker, 1994). These findings suggest that future studies should quantify visceral afferent sensitivity, as it may significantly contribute to nausea and represent an unmeasured source of individual variability in motion sickness susceptibility (Minoretti et al., 2024; Talsma et de Winkel, 2025).

In conclusion, while the symptomatology of motion sickness induced by parabolic flight (PF) and virtual reality (VR) largely overlaps, a key distinction emerged: systolic blood pressure decreased exclusively in PF. Among the physiological markers assessed, skin greenness proved the most reliable and measurable parameter across both paradigms. However, individual susceptibility to motion sickness remains challenging to predict, even when baseline physiological data are considered, and gastrointestinal symptoms—the most debilitating—remained elusive to objective prediction. These results highlight the complexity of motion sickness physiology and emphasize the need to account for paradigm-specific mechanisms in future research.

## AUTHORS CONTRIBUTIONS

*T.B. and E.G.*: Conceived and designed the study, performed experiments, analyzed data, conducted statistical analyses, interpreted results and wrote manuscript.

*E.D.*: Performed experiments

*J.C.:* Contributed to results interpretation and assisted with manuscript revisions C.M.: Contributed to data analysis

*D.G.*: Assisted in project conception and participant inclusion visit

---

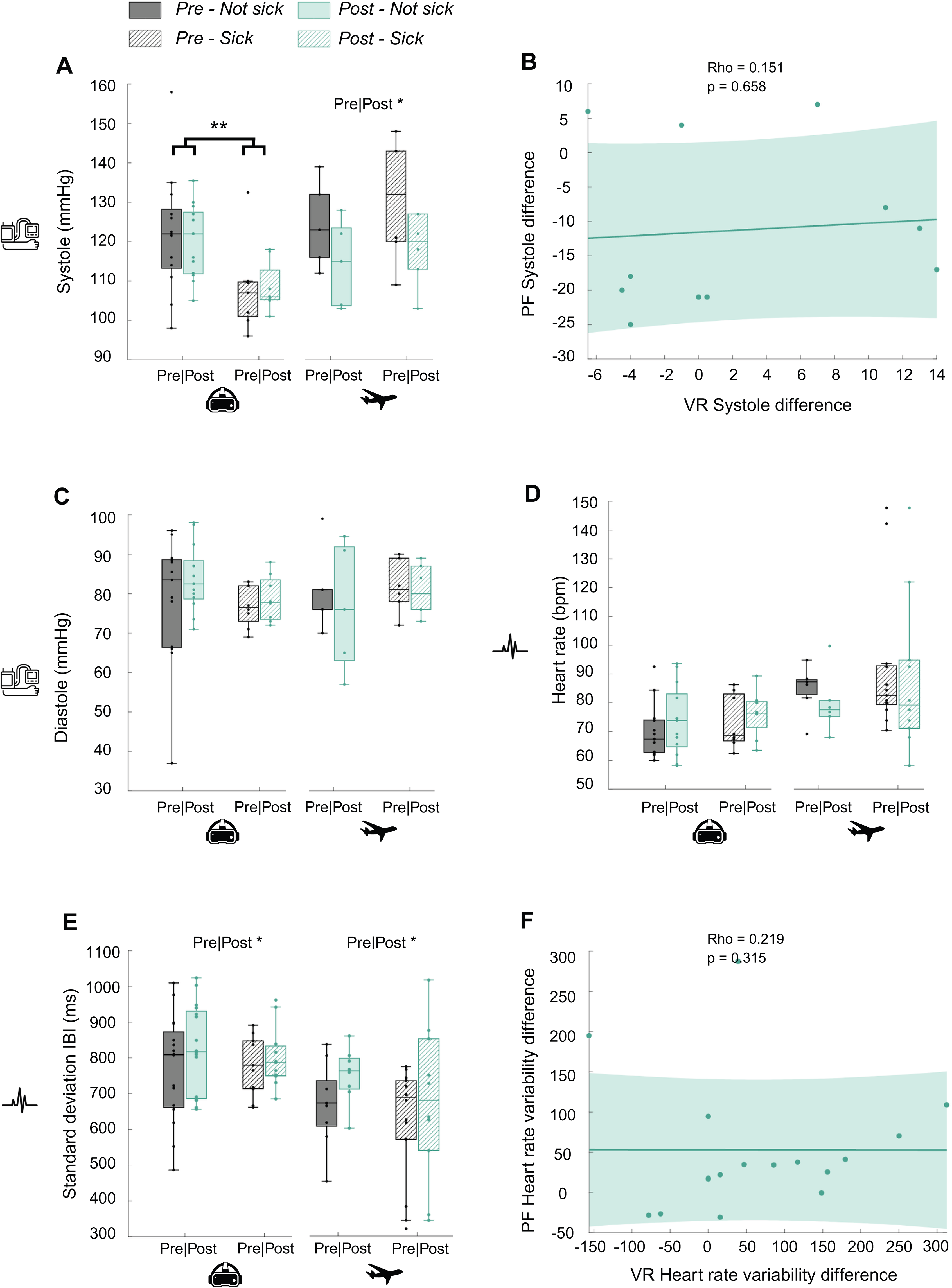

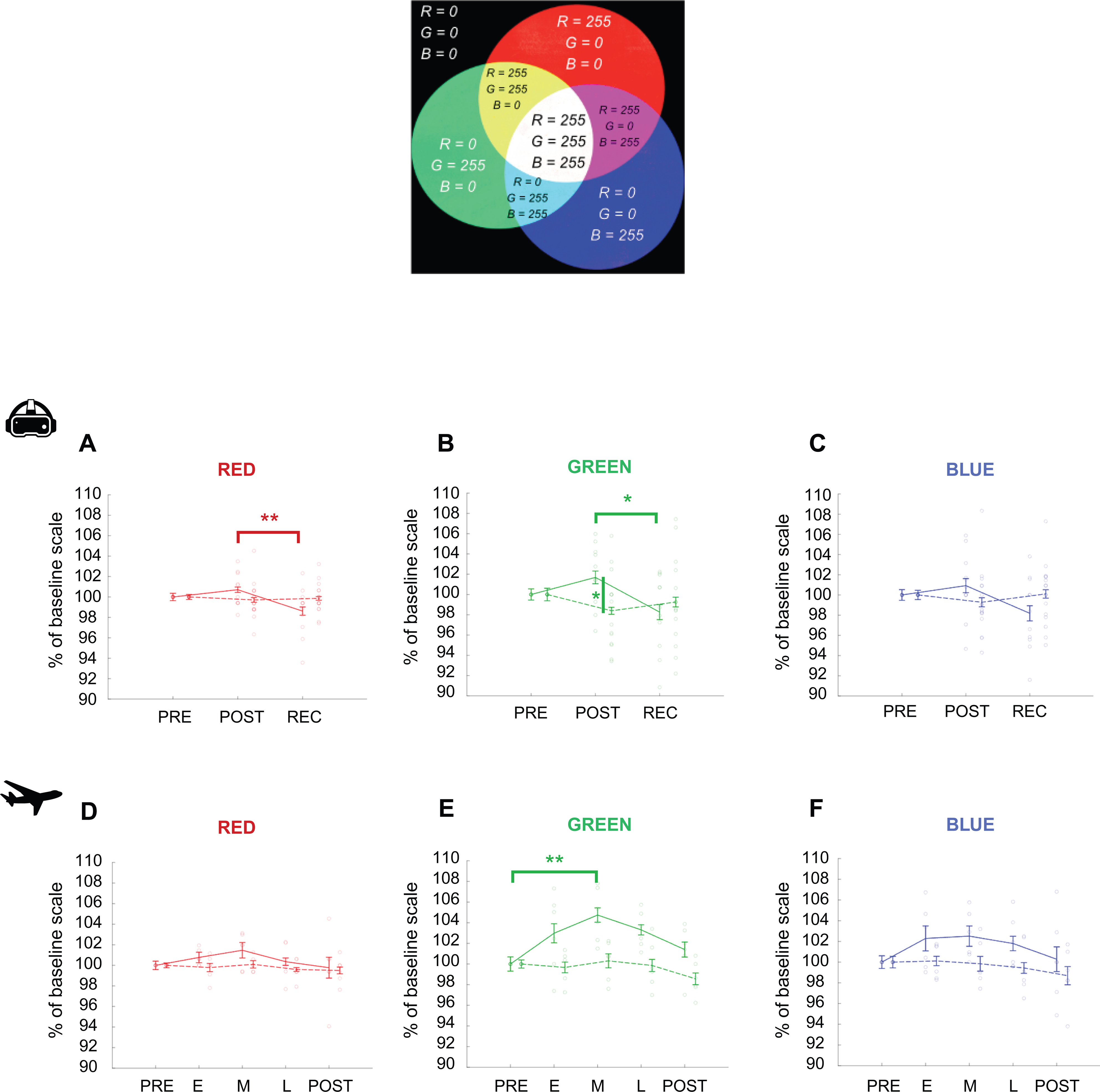

## REFERENCES

Altena, E. et al. How sleep problems contribute to simulator sickness: Preliminary results from a realistic driving scenario. Journal of Sleep Research 28, e12677 (2019).

Arnold, J. T., O’Keeffe, K., McDaniel, C., Hodder, S. & Lloyd, A. Effect of virtual reality and whole-body heating on motion sickness severity: A combined and individual stressors approach. Displays 60, 18–23 (2019).

Babic, T. & Browning, K. N. The role of vagal neurocircuits in the regulation of nausea and vomiting. Eur J Pharmacol 722, 38–47 (2014).

Bestaven, E., Kambrun, C., Guehl, D., Cazalets, J.-R. & Guillaud, E. The influence of scopolamine on motor control and attentional processes. PeerJ 4, e2008 (2016).

Bonnard, T., Doat, E., Cazalets, J., Guehl, D. & Guillaud, E. Visual and vestibular reweighting after cyber- and space-sickness. Experimental Physiology 111, 240–256 (2026).

Bonnard, T., Araka, L., Smith Reina, A., van Berkel, J., Morvan, C., Morgat, C., Guehl, D., Guillaud, E. & Altena, E. Sleep’s protective role against space sickness. (submitted)

Bonnard, T., Morello, A., Doat, E., Cazalets, J-R., Guehl, D. & Guillaud, E. Resilience of otolithic reflex to gravitational changes. NPJ Microgravity (minor reviews)

Borison, H. L. & Wang, S. C. PHYSIOLOGY AND PHARMACOLOGY OF VOMITING. Pharmacological Reviews 5, 193–230 (1953).

Choukèr, A., et al. Motion Sickness, Stress and the Endocannabinoid System. PLoS ONE 5, e10752 (2010).

Clément, G., Macaulay, T. R., Moudy, S. C., Kuldavletova, O. & Wood, S. J. Back to the future—revisiting Skylab data on ocular counter-rolling and motion sickness. Front. Physiol. 14, 1303938 (2023).

Clément, G. R. et al. Challenges to the central nervous system during human spaceflight missions to Mars. Journal of Neurophysiology 123, 2037–2063 (2020).

Clément, G. & Reschke, M. F. Relationship between motion sickness susceptibility and vestibulo-ocular reflex gain and phase. VES 28, 295–304 (2018).

Cohen, B., Dai, M., Yakushin, S. B. & Cho, C. The neural basis of motion sickness. Journal of Neurophysiology 121, 973–982 (2019).

Czeisler, M. É. et al. Validation of the motion sickness severity scale: Secondary analysis of a randomized, double-blind, placebo-controlled study of a treatment for motion sickness. PLoS ONE 18, e0280058 (2023).

Dahlman, J., Sjörs, A., Lindström, J., Ledin, T. & Falkmer, T. Performance and Autonomic Responses During Motion Sickness. Hum Factors 51, 56–66 (2009).

Daviaux, Y. et al. Event-Related Electrodermal Response to Stress: Results From a Realistic Driving Simulator Scenario. Hum Factors 62, 138–151 (2020).

Dijk, D.-J. et al. Amplitude Reduction and Phase Shifts of Melatonin, Cortisol and Other Circadian Rhythms after a Gradual Advance of Sleep and Light Exposure in Humans. PLoS ONE 7, e30037 (2012).

Dobie, T. G. Motion Sickness: A Motion Adaptation Syndrome. vol. 6 (Springer International Publishing, Cham, 2019).

Eiken, O., Tipton, M. J., Kölegard, R., Lindborg, B. & Mekjavic, I. B. Motion sickness decreases arterial pressure and therefore acceleration tolerance. Aviat Space Environ Med 76, 541–546 (2005).

Gallagher, M. & Ferrè, E. R. Cybersickness: a Multisensory Integration Perspective. Multisens. Res. 31, 645–674 (2018).

Golding, J. F. Motion sickness susceptibility. Autonomic Neuroscience 129, 67–76 (2006).

Golding, J. F., Paillard, A. C., Normand, H., Besnard, S. & Denise, P. Prevalence, Predictors, and Prevention of Motion Sickness in Zero-G Parabolic Flights. aerosp med hum perform 88, 3–9 (2017).

Golding, J. F., Rafiq, A. & Keshavarz, B. Predicting Individual Susceptibility to Visually Induced Motion Sickness by Questionnaire. Front. Virtual Real. 2, 576871 (2021).

Harm, D. L. & Schlegel, T. T. Predicting motion sickness during parabolic flight. Autonomic Neuroscience 97, 116–121 (2002).

Heer, M. & Paloski, W. H. Space motion sickness: Incidence, etiology, and countermeasures. Autonomic Neuroscience 129, 77–79 (2006).

Holmes, S. R., King, S., Rollin Scott, J. R. & Clemes, S. Facial Skin Pallor Increases During Motion Sickness. Journal of Psychophysiology 16, 150–157 (2002).

Homick, J. L. Space motion sickness. Acta Astronautica 6, 1259–1272 (1979).

2009 36th Annual Computers in Cardiology Conference. (I E E E, Place of publication not identified, 2009).

Javaid, A., Chouhna, H., Varghese, B., Hammam, E. & Macefield, V. G. Changes in skin blood flow, respiration and blood pressure in participants reporting motion sickness during sinusoidal galvanic vestibular stimulation. Experimental Physiology 104, 1622–1629 (2019).

Kaplan, J. et al. The influence of sleep deprivation and oscillating motion on sleepiness, motion sickness, and cognitive and motor performance. Autonomic Neuroscience 202, 86–96 (2017).

Kennedy, R. S., Lane, N. E., Berbaum, K. S. & Lilienthal, M. G. Simulator Sickness Questionnaire: An Enhanced Method for Quantifying Simulator Sickness. The International Journal of Aviation Psychology 3, 203–220 (1993).

Keshavarz, B., Peck, K., Rezaei, S. & Taati, B. Detecting and predicting visually induced motion sickness with physiological measures in combination with machine learning techniques. International Journal of Psychophysiology 176, 14–26 (2022).

Kolev, O. I., Möller, C., Nilsson, G. & Tibbling, L. Responses in Skin Microcirculation to Vestibular Stimulation Before and During Motion Sickness. Can. j. neurol. sci. 24, 53–57 (1997).

Kourtesis, P., Amir, R., Linnell, J., Argelaguet, F. & MacPherson, S. E. Cybersickness, Cognition, & Motor Skills: The Effects of Music, Gender, and Gaming Experience. IEEE Trans. Visual. Comput. Graphics 29, 2326–2336 (2023).

Lackner, J. R. Motion sickness: more than nausea and vomiting. Exp Brain Res 232, 2493–2510 (2014).

Lentz, J. M. The Aerospace Medical Panel Symposium on Motion Sickness: Mechanisms, Prediction, Prev∼ention, and Treatment Held at Williamsburg, Virginia on 3–4 May 1984. https://apps.dtic.mil/sti/html/tr/ADP004643/index.html (1984).

Leung, A. K. & Hon, K. L. Motion sickness: an overview. DIC 8, 1–11 (2019).

Li, C. et al. Multi-Dimensional and Objective Assessment of Motion Sickness Susceptibility Based on Machine Learning. Front. Neurol. 13, 824670 (2022).

Liu, Y., Wang, L., Liu, J. & Di, Y. A study of human skin and surface temperatures in stable and unstable thermal environments. Journal of Thermal Biology 38, 440–448 (2013).

Reyero Lobo, P. & Perez, P. Heart Rate Variability for Non-Intrusive Cybersickness Detection. in *ACM International Conference on Interactive Media Experiences* 221–228 (ACM, Aveiro JB Portugal, 2022). doi:10.1145/3505284.3529973.

Malik, M. Heart rate variability: Standards of measurement, physiological interpretation, and clinical use. Circulation 93, 1043–1065 (1996).

Mazdeyasna, S., Ghassemi, P. & Wang, Q. Best Practices for Body Temperature Measurement with Infrared Thermography: External Factors Affecting Accuracy. Sensors 23, 8011 (2023).

Metzulat, M., Metz, B., Landau, A., Neukum, A. & Kunde, W. Too sick to take over? − Impact of car sickness on cognitive performance related to driving in the context of automated driving. Transportation Research Part F: Traffic Psychology and Behaviour 109, 480–500 (2025).

Miller, A. D. & Leslie, R. A. The Area Postrema and Vomiting. Frontiers in Neuroendocrinology 15, 301–320 (1994).

Minoretti, P., Fortuna, G., Lavdas, K. & D’Acquino, D. Potential Biomarkers of Resilience to Microgravity Hazards in Astronauts. Cureus 16, e57173 (2024).

Mittelstaedt, H. Somatic graviception. Biological Psychology 42, 53–74 (1996).

Nalivaiko, E., Rudd, J. A. & So, R. H. Motion sickness, nausea and thermoregulation: The “toxic” hypothesis. Temperature 1, 164–171 (2014).

Nobel, G., Tribukait, A., Mekjavic, I. B. & Eiken, O. Effects of motion sickness on thermoregulatory responses in a thermoneutral air environment. Eur J Appl Physiol 112, 1717–1723 (2012).

Oman, C. M. & Cullen, K. E. Brainstem processing of vestibular sensory exafference: implications for motion sickness etiology. Exp Brain Res 232, 2483–2492 (2014).

Ortega, H. J. & Harm, D. L. Space and Entry Motion Sickness. in Principles of Clinical Medicine for Space Flight (eds Barratt, M. R. & Pool, S. L.) 211–222 (Springer New York, New York, NY, 2008). doi:10.1007/978-0-387-68164-1_10.

Ortonne, J.-P. La couleur de la peau normale ou anormale. Annales de Dermatologie et de Vénéréologie 139, S73–S77 (2012).

Otto, B., Riepl, R. L., Klosterhalfen, S. & Enck, P. Endocrine correlates of acute nausea and vomiting. Autonomic Neuroscience 129, 17–21 (2006).

Reason, J. T. & Brand, J. J. Motion Sickness. vol. 31 (1975).

Reschke, M. F., Good, E. F. & Clément, G. R. Neurovestibular Symptoms in Astronauts Immediately after Space Shuttle and International Space Station Missions. OTO Open 1, 2473974X17738767 (2017).

Schmidt, E. A., Kuiper, O. X., Wolter, S., Diels, C. & Bos, J. E. An international survey on the incidence and modulating factors of carsickness. Transportation Research Part F: Traffic Psychology and Behaviour 71, 76–87 (2020).

Séba, M.-P., Maillot, P., Hanneton, S. & Dietrich, G. Influence of Normal Aging and Multisensory Data Fusion on Cybersickness and Postural Adaptation in Immersive Virtual Reality. Sensors 23, 9414 (2023).

Stern, R. M., Koch, K. L., & Andrews, P. L. R. (2011). Nausea: Mechanisms and management. Oxford University Press.

Tal, D. et al. Artificial Horizon Effects on Motion Sickness and Performance. Otology & Neurotology 33, 878–885 (2012).

Talsma, T. M. W. & De Winkel, K. N. The gut feeling in motion sickness. Commun Biol 8, 1497 (2025).

von Gierke, H. E. & Parker, D. E. Differences in otolith and abdominal viscera graviceptor dynamics: implications for motion sickness and perceived body position. Aviat Space Environ Med 65, 747–751 (1994).

Wan, H., Hu, S. & Wang, J. Correlation of Phasic and Tonic Skin-Conductance Responses with Severity of Motion Sickness Induced by Viewing an Optokinetic Rotating Drum. Percept Mot Skills 97, 1051–1057 (2003).

Weerts, A. P. et al. Intranasal scopolamine affects the semicircular canals centrally and peripherally. Journal of Applied Physiology 119, 213–218 (2015).

Yates, B. J., Bolton, P. S. & Macefield, V. G. Vestibulo-Sympathetic Responses. in Comprehensive Physiology (ed. Terjung, R.) 851–887 (Wiley, 2014). doi:10.1002/cphy.c130041.

Zhang, L. et al. Motion Sickness: Current Knowledge and Recent Advance. CNS Neurosci Ther 22, 15–24 (2016).

Pathology of emesis. in Handbook of Clinical Neurology vol. 117 337–352 (Elsevier, 2013).

Thermoregulation and nausea. in Handbook of Clinical Neurology vol. 156 445–456 (Elsevier, 2018).

